# Role of DCP1-DCP2 complex regulated by viral and host microRNAs in DNA virus infection

**DOI:** 10.1101/341362

**Authors:** Yuechao Sun, Xiaobo Zhang

## Abstract

The DCP1-DCP2 complex can regulate the animal antiviral immunity by the decapping of retrovirus RNAs and the suppression of RNAi pathway. However, the influence of DCP1-DCP2 complex on DNA virus infection and the regulation of DCP1-DCP2 complex by microRNAs (miRNAs) remain unclear. In this study, we investigated the role of miRNA-regulated DCP1-DCP2 complex in DNA virus infection. Our results suggested that the DCP1-DCP2 complex played a positive role in the infection of white spot syndrome virus (WSSV), a DNA virus of shrimp. The N-terminal regulatory domain of DCP2 was interacted with the EVH1 domain of DCP1, forming the DCP1-DCP2 complex. Furthermore, a host shrimp miRNA (miR-87) inhibited WSSV infection by targeting the host DCP2 gene and a viral miRNA (WSSV-miR-N46) took a negative effect on WSSV replication by targeting the host DCP1 gene. Therefore, our study provided novel insights into the underlying mechanism of DCP1-DCP2 complex and its regulation by miRNAs in virus-host interactions.

The DCP1-DCP2 complex can regulate the animal antiviral immunity by the decapping of retrovirus RNAs and the suppression of RNAi pathway. In the present study, the findings indicated that the silencing of the DCP1-DCP2 complex inhibited the infection of WSSV, a DNA virus of shrimp, suggesting that the DCP1-DCP2 complex facilitated DNA virus infection. Due to the suppressive role of the DCP1-DCP2 complex in RNAi pathway against virus infection, the DCP1-DCP2 complex could promote WSSV infection in shrimp. In this context, our study contributed a novel aspect of the DCP1-DCP2 complex in virus-host interactions. Our study revealed that the host and viral miRNAs could regulate the DCP1-DCP2 complex to affect virus infection. Therefore, our study provided novel insights into the miRNA-mediated regulation of DCP1-DCP2 complex took great effects on RNAi immunity of invertebrates against virus infection.

## Introduction

Classical virus infection in the host cell is initiated by interactions between viral capsid or envelope proteins and host cell surface receptors. The internalization of virions is either through the fusion of the viral envelope and the host plasma membrane, or through the endocytosis pathway, causing the virions to escape from the endocytosa or other small vesicles and enter the cytoplasm (1). Cell receptors attached to the host can directly trigger conformational changes in the surface structure of the virus or activate specific signaling pathways that facilitate the entry of the virus (1). The life cycle of a virus begins with the entry of the host cell. Replication of viral genomes, synthesis of viral proteins, assembly of viral particles, and release of viruses from host cells depend largely on host mechanisms (1, 2). It is reported that the stability of viral mrna is regulated by the dcp1-dcp2 complex located in the P-body (the processing bodies) (3, 4). The DCP1-DCP2 complex can trigger mRNA decapping. DCP2 catalyzed dissection releases m7G and a single 5’ phosphorylated mRNA. This is considered to be an irreversible process, and the target is that mRNA is degraded by exonuclease Xrn1, 5’ to 3’ (5). DCP2 protein contains n-terminal Nudix/MutT motifs, which are usually present in pyrophosphatase and are essential for decapping (4, 6). Except for the DCP2-DCP1 complex, Pat1 (Decapping activator and translation repressor) (7–10), Dhh1 (Decapping activator and translation repressor) (10–13) and the Lsm1-7 complex (Decapping activator) (10, 12, 14) are involved in the decapping of mRNAs. At present, the decapping of retrovirus RNAs by the DCP2-DCP1 complex has been well characterized (3, 4). However, the role of DCP1-DCP2 complex in DNA virus infection remains unclear.

Although the DCP1-DCP2 complex affects the mRNA stability, the regulation of DCP1-DCP2 complex mediated by microRNAs (miRNAs) has not been extensively explored. During many eukaryotic cellular processes the miRNA pathway is essential, in especial as virus-host interaction, development, apoptosis, immune response, tumorigenesis and homeostasis (15–17). Primary miRNAs (pri-miRNAs) and precursor-miRNAs (pre-miRNAs) are the essential steps of miRNAs in cell nucleus (18–20). After being transported into cytoplasm, pre-miRNAs are processed by Dicer, producing ~ 22 bp mature miRNA duplexes. The RNA-induced silencing complex (RISC) is formed after the loading of the guiding strand of miRNA onto Argonaute (Ago) protein (18). The target mRNA is bound to the miRNA and then it will be cleaved by the Ago protein, in the RISC (18). Recently, it has been reported that phosphorylation and dephosphorylation of Ago2 protein in human body have a significant impact on the role of miRNA in RISC. (15). In the virus-host interactions, the gene expressions can be regulated by host and/or virus miRNAs (20–31). In shrimp, the host miRNAs expression are altered by the infection of white spot syndrome virus (WSSV), a virus with a double-stranded DNA genome (25, 27, 30, 31). Shrimp miR-7 can target the WSSV early gene wsv477, thus inhibiting virus infection (17), while a viral miRNA can target the shrimp caspase 8 gene to suppress the host antiviral apoptosis (25). It has been reported that virus-originated mirnas promote viral latency during viral infection through RNA editing (32). At present, however, the influence of miRNA-mediated regulation of the DCP1-DCP2 complex on virus infection remains to be investigated.

To address the influence of DCP1-DCP2 complex on DNA virus infection and the role of the miRNA-regulated DCP1-DCP2 complex in virus infection, shrimp and WSSV miRNAs targeting the DCP1-DCP2 complex were characterized in this study. The results indicated that shrimp miR-87 and viral WSSV-miR-N46 (a viral miRNA) could suppress virus infection by targeting the DCP1-DCP2 complex.

## Materials and methods

### Shrimp culture and WSSV challenge

Shrimp (*Marsupenaeus japonicus*), 10 to 12 cm in length, were cultured in groups of 20 individuals in the tank filled with seawater at 25°C (23). To ensure that shrimp were virus-free before experiments, PCR using WSSV-specific primers (5’-TATTGT CTC TCCTGACGTAC-3’ and 5’-CACATTCTTCACGAGTCTAC-3’) was performed to detect WSSV in shrimp (23). The virus-free shrimp were infected with WSSV inoculum (10^5^ copies/ml) by injection at 100 μl/shrimp into the lateral area of the fourth abdominal segment of shrimp (23). At different time postinfection, three shrimp were randomly collected for each treatment. The shrimp hemocytes were collected for later use.

### Analysis of WSSV copies with quantitative real-time PCR

The genomic DNA of WSSV was extracted with a SQ tissue DNA kit (Omega Bio-tek, Norcross, GA, USA) according to the manufacturer’s instruction. The extracted DNA was analyzed by quantitative real-time PCR with WSSV-specific primers and WSSV-specific TaqMan probe (5’-FAM-TGCTGCCGTCTCCAA-TAMRA-3’) as described previously (Huang et al, 2014) (23). The PCR procedure was 95°C for 1 min, followed by 40 cycles of 95°C for 30 s, 52°C of 30 s, and 72°C for 30 s (23).

### Detection of mRNA or miRNA by Northern blotting

The RNA was extracted from shrimp hemocytes with mirVana miRNA isolation kit (Ambion, USA). After separation on a denaturing 15% polyacrylamide gel containing 7M urea, the RNA was transferred to a Hybond-N+ nylon membrane, followed by ultraviolet cross-linking (23). The membrane was prehybridized in DIG (digoxigenin) Easy Hyb granule buffer (Roche, Basel, Switzerland) for 0.5 h at 42°C and then hybridized with DIG-labeled miR-87 (5’-GAGGGGAAAAGCCATACGCT TA-3’), WSSV-miR-N46 (5’-AGUGCCAAGAUAACGGUUGAAG-3’), U6 (5’-GG GCCATGCTAATCTTCTCTGTATCGTT-3’), wsv477 (5’-CGAT TTCGGCAGGC CAGTTGTCAGA-3’), DCP2 (5’-CCAGAAACCCTGAACTAAGAGAA-3’) or actin (5’-CTCGCTCGGCGGTGGTCGTGAAGG-3’) probe at 42 °C overnight (23). Subsequently the detection was performed with the DIG High Prime DNA labeling and detection starter kit II (Roche).

### Silencing or overexpression of miR-87 or WSSV-miR-N46 in shrimp

To knock down miR-87 or WSSV-miR-N46, an anti-miRNA oligonucleotide (AMO) was injected into WSSV-infected shrimp (23). AMO-miR-87 (5’-TGTACGTTTC TGGAGC-3’) and AMO-WSSV-miR-N46 (5’-CTTCAACCGTTATCTTGGCACT-3’) were synthesized (Sangon Biotech, Shanghai, China) with a phosphorothioate backbone and a 2’-O-methyl modification at the 12th nucleotide. AMO (10 nM) and WSSV (10^5^ copies/ml) were co-injected into virus-free shrimp at a 100 μl/shrimp (23). At 16 h after the co-injection, AMO (10 nM) was injected into the same shrimp. As controls, AMO-miR-87-scrambled (5’-TTGCATGTCTGTCGAG-3’), AMO-WSSV-miR-N46-scrambled (5’-TTGCATGTCTGTCGAG-3’), WSSV alone (10^5^ copies/ml) and phosphate buffered saline (PBS) were included in the injections. To overexpress miR-87 or WSSV-miR-N46, the synthesized miR-87 (5’-TAAGCGTAT GGCTTTTCCCCTC-3’) (10 nM) or WSSV-miR-N46 (5’-AGTGCCAAGATAACG GTTGAAG-3’) and WSSV (10^5^ copies/ml) were co-injected into shrimp. As controls, miR-87-scrambled (5’-TATCGCATAGGCTTTTCCCCTC-3’), WSSV-miR-N46-scrambled (5’-ATTTGACAGATGCCTAGTACCAG-3’), WSSV alone (10^5^ copies/ml) and PBS were used (23). The miRNAs were synthesized by Sangon Biotech (Shanghai, China).

At different time after treatment with AMO or miRNA, three shrimp were collected at random for each treatment. The shrimp hemocytes were collected for later use. At the same time, the cumulative mortality of shrimp was examined daily. All the experiments were biologically repeated three times.

### Prediction of miRNA target genes

To predict the target genes of a miRNA, four independent computational algorithms including TargetScan 5.1 (http://www.targetscan.org), miRanda (http://www.microrna.org/), Pictar (http://www.pictar.mdc-berlin.de/) and miRlnspector (http://www.Imbb.Forth.gr/microinspector) were used (23). The overlapped genes predicted by the four algorithms were the potential targets of the miRNA.

### Cell culture, transfection and fluorescence assays

Insect High Five cells (Invitrogen, USA) were cultured with Express Five serum-free medium (Invitrogen) containing L-glutamine (Invitrogen) at 27°C (23). To determine the dosage of a synthesized miRNA, 10, 50, 100, 200, 500 or 1000 pM of miRNA was transfected into cells (23). Then the miRNA expression in cells was detected with quantitative real-time PCR. It was indicated that the transfection of miRNA at 100 pM or more could overexpress miRNA in cells. The insect cells were co-transfected with EGFP, EGFP-DCP2-3’UTR, EGFP-ΔDCP2-3’UTR, EGFP-DCP1-3’UTR or EGFP-ΔDCP1-3’UTR and miRNA (miR-87 or WSSV-miR-N46). All the miRNAs were synthesized by Shanghai GenePharma Co., Ltd (Shanghai, China). At 48h after co-transfection, the fluorescence intensity of cells was evaluated with a Flex Station II microplate reader (Molecular Devices, USA) at 490/510 nm excitation/emission (Ex/Em) (23). The experiments were biologically repeated three times.

### Western blot analysis

Shrimp tissues were homogenized with a lysis buffer (50 mM Tris-HCl, 150 mM NaCl, 0.1% SDS, 1% Triton X-100, 1 mM phenylmethylsulfonyl fluoride, pH7.8) and then centrifuged at 10,000×g for 10 min at 4°C. The proteins were separated by 12.5% SDS-polyacrylamide gel electrophoresis and then transferred onto a nitrocellulose membrane. The membrane was blocked with 5% non-fat milk in TBST (10 mM Tris-HCl, 150 mM NaCl, 20% Tween 20, pH7.5) for 2 h at room temperature, followed by incubation overnight with a primary antibody. The antibodies were prepared in our laboratory. After washes with TBST, the membrane was incubated with horseradish peroxidase-conjugated secondary antibody (Bio-Rad, USA) for 2 h at room temperature. Subsequently the membrane was detected using a Western Lightning Plus-ECL kit (Perkin Elmer, USA).

### RNAi (RNA interference) assay in shrimp

To silence gene expression in shrimp, RNAi assay was conducted. The small interfering RNA (siRNA) specifically targeting the *DCP1* or *DCP2* gene was designed with the 3’ UTR of the *DCP1* or *DCP2* gene, generating DCP1-siRNA (5’-A AUCGCAGUUGCUAUGCGUUGGACG-3’) or DCP2-siRNA (5’-GCGGAAGAC CGUGCCCGUAAUAUAA-3’). As a control, the sequence of DCP1-siRNA or DCP2-siRNA was randomly scrambled (DCP1-siRNA-scrambled, 5’-GACAUUAAG AUAUAUAUGG-3’; DCP2-siRNA-scrambled, 5’-CGCCUUCUGGCACGGGCAUUAUAUU-3’). All the siRNAs were synthesized by the in vitro transcription T7 kit (TaKaRa, Japan) according to the manufacturer’s instructions. The synthesized siRNAs were quantified by spectrophotometry. The shrimp were co-injected with WSSV (10^4^ copies/shrimp) and siRNA (4nM). PBS and WSSV alone (10^4^ copies/shrimp) were included in the injections as controls (23). At 0, 24, 36 and 48 h after infection, the hemocytes of three shrimp, randomly selected from each treatment, were collected for later use. At the same time, the cumulative mortality of shrimp was examined daily (23). All the experiments were biologically repeated three times.

### Co-immunoprecipitation

Shrimp hemocytes were lysed with ice-cold cell lysis buffer (Beyotime). Then the lysate was incubated with Protein G-agarose beads (Invitrogen, Carlsbad, CA, USA) for 2h at room temperature, followed by incubation with DCP2-specific antibody overnight at 4^0^C. After washes three times with ice-cold lysis buffer, the immuno-complex was subjected to SDS-PAGE with Coomassie blue staining. The proteins were identified with mass spectrometry using a Reflex IV MALDI-TOF mass spectrometer (Bruker Daltonik, Bremen, USA). The spectra were processed by the Xmass software (Bruker Daltonik, Bremen) and the peak lists of the mass spectra were used for peptide mass fingerprint analyses with the Mascot software (Matrix Science).

### Cloning of full-length cDNAs of shrimp *DCP1* and *DCP2* gene

The full-length *DCP1* and *DCP2* cDNAs were obtained by rapid amplification of cDNA ends (RACE) using a 5’/3’ RACE kit (Roche, Indianapolis, IN, USA). RACEs were conducted according to the manufacturer’s instructions using DCP1-specific primers (5’RACE, 5’-CCTGGGACACTTGAAG-3’ and 5’-GGGTAAACCAGTGCC-3’; 3’RACE, 5’-GCCCCACAGTCCCACCCACCT-3’ and 5’-CCCAGGAGGAGCA CCAATCTCA-3’) or DCP2-specific primers (5’RACE, 5’-GGGAACCATTTCAGT TGCT-3’ and 5’-GCCAGAAACCCTGAACTAAG-3’; 3’RACE, 5’-ATTGGAGAGC AGTTTGTGAGAC-3’ and 5’-TTTACATCATCCCAGGCG-3’). PCR products were cloned into pMD-19 vector (Takara, Japan) and sequenced.

### Interactions between DCP1 and DCP2 domains

To explore the interaction between DCP1 and DCP2 proteins, the full-length and domain deletion mutants of DCP1 and DCP2 were cloned into pIZ/EGFP V5-FLAG and pIZ/EGFP V5-His (Invitrogen, USA), respectively. The full-length and deletion mutants of DCP1 and DCP2 were amplified by PCR with sequence-specific primers (full-length DCP1, 5’-GGAAGATCTATGCGCTAAGGTTTTATTTGGAAAAA-3’ and 5’-CCGCTCGAGTGACTTATCGTCGTCATCCTTGTAATCCAAACAACCTTTGATAGAGAGAT-3’; DCP1 EVH1 domain, 5’-GGAAGATCTATGCGCTAAGGTTTTATTTGGAAAAA-3’ and 5’-CCGCTCGAGTGACTTATCGTCGTCATCCTTGTAATCTATGTCCCCTCCAGGTGCCCCA-3’; DCP1 C-terminal extension region, 5’-GGAAGATCTGAATGACAAATCAAGTGA-3’ and 5’-CCGCTCGAG TGACTTATCGTCGTCATCCTTGTAATCCAAACAACCTTTGATAGAGAGAT-3’; full-length DCP2, 5’-GGAAGATCTATGGCCCCACCAACAGGTGGAAAA-3’ and 5’-TCCCCGCGGTTAATGGTGATGGTGATGATGCCAAGACAGCATCACATCGGCCC-3’; DCP2 N-regulatory domain, 5’-CGCGGATCCGATGAAGAACCACATTGTTGTGCC-3’ and 5’-TCCCCGCGGTTAATGGTGATGGTGATGATGCCAAGACAGCATCACATCGGCCC-3’; DCP2 C-terminal divergent region, 5’-CGCGGATCCGATGAAGAACCACATTGTTGTGCC-3’ and 5’-TCCCCGCGG TTAATGGTGATGGTGATGATGCTGGCGGTCAGAGGTACTGGTG-3’; DCP2 Nudix domain, 5’-CG CGGATCCATGGCCCCACCAACAGGTGGAAAA-3’ and 5’-TCCCCGCGGCACATTAATTTTCCCTTTTGG-3’, and 5’-TCCCCGCGGATGG CCCCACCAACAGGTGGAAAA-3’ and 5’-TCCCCGCGGTTAATGGTGATGGTG ATGA TGCCAAGA CAGCATCACATCGGCCC-3’).

The constructs were co-transfected into insect High Five cells at 70% confluence using Cellfectin transfection reagent (Invitrogen, USA) according to the manufacturer’s protocol. The cells were cultured at 27 °C in Express Five serum-free medium (Invitrogen) supplemented with L-glutamine (Invitrogen). At 48 h after co-transfection, the cells were subjected to immunoprecipitation assays with anti-His or anti-FLAG antibody, followed by Western blot analysis.

### Statistical analysis

All the numerical data presented were analyzed by one-way analysis of variance (ANOVA) to calculate the means and standard deviations of triplicate assays.

## Results

### Role of shrimp DCP2 in virus infection

To characterize the role of shrimp DCP2 in virus infection, the expression level of DCP2 was examined in shrimp in response to WSSV infection. The results indicated that DCP2 was significantly upregulated in virus-challenged shrimp (Fig 1A), suggesting that DCP2 played an important role in virus infection.

**Fig 1.**
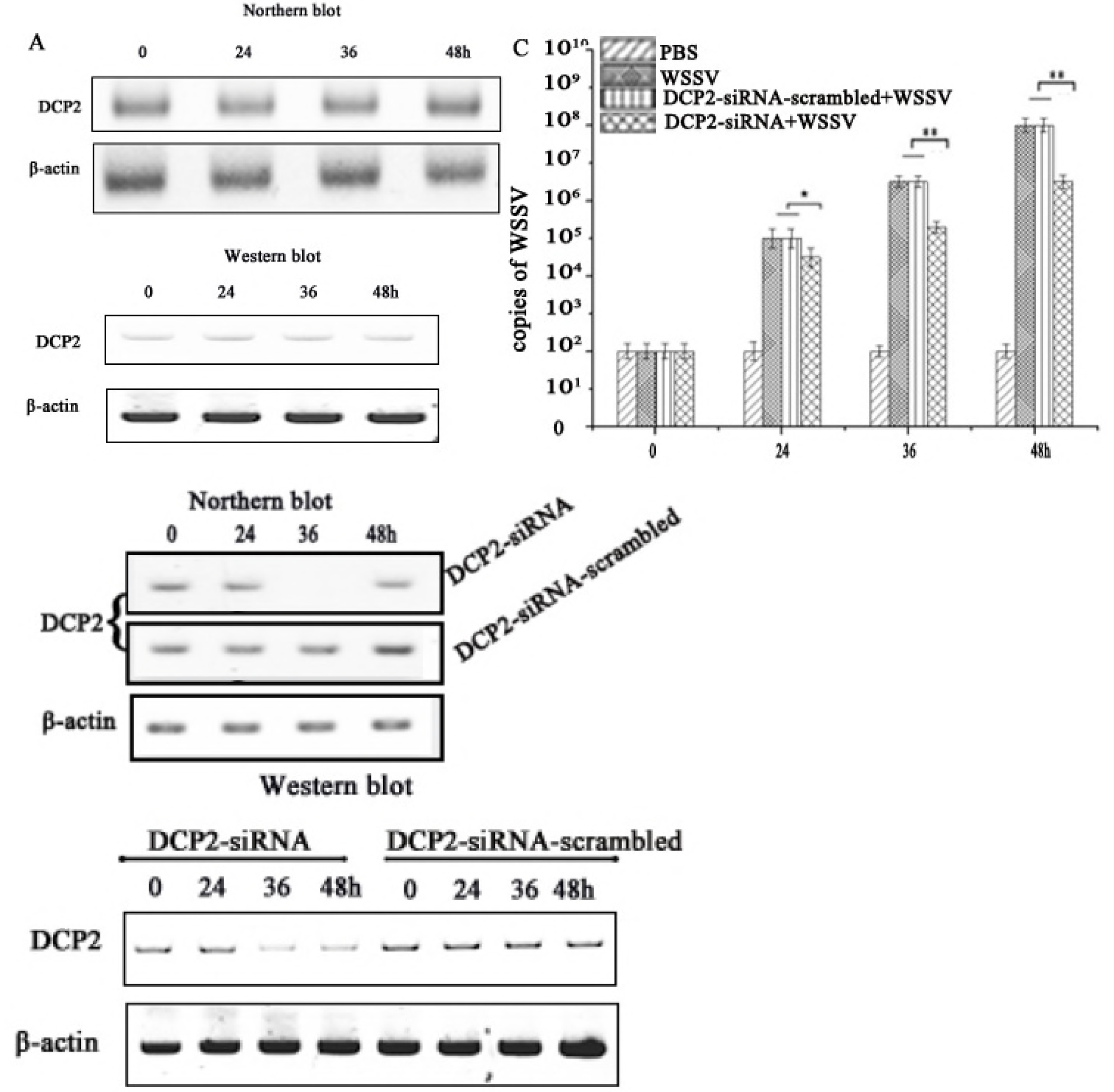
Role of shrimp DCP2 in virus infection. (A) Expression level of DCP2 in shrimp in response to virus infection. Shrimp were challenged with WSSV. At different times post-infection, the expression level of DCP2 in shrimp hemocytes was examined by Northern blotting or Western blotting. Shrimp β-actin was used as a control. Numbers indicated the time post-infection. Probes or antibodies used were shown on the left. (B) Knockdown of DCP2 by siRNA in shrimp. Shrimp were injected with DCP2-siRNA to silence DCP2 expression. As a control, DCP2-siRNA-scrambled was included in the injection. At different time after injection, the *DCP2* mRNA and protein levels were examined by Northern blot and Western blot, respectively. Actin was used as a control. The probes or antibodies were indicated on the left. (C) Influence of DCP2 silencing on virus infection in shrimp. Shrimp were co-injected with DCP2-siRNA and WSSV. At different time post-infection, the WSSV copies were examined with quantitative real-time PCR (*, *p*< 0.05; **, *p*< 0.01).

To explore the influence of DCP2 silencing on virus infection, the DCP2 expression was knocked down by sequence-specific DCP2-siRNA in shrimp (Fig 1B). The results revealed that the DCP2 silencing resulted in significant decreases of WSSV copies compared with the controls (Fig 1C), showing that DCP2 played an essential role in WSSV infection.

### Proteins interacted with DCP2

To elucidate the mechanism of DCP2-mediated antiviral immunity in shrimp, the proteins interacted with DCP2 were characterized. The results of co-immunoprecipitation assays using shrimp DCP2-specific antibody indicated that two proteins were obtained compared with the control (Fig 2A). Mass spectrometry identification revealed that the two proteins were DCP1 and DCP2 (Fig 2A). These data showed that DCP2 was interacted with DCP1 in shrimp. To confirm the interaction between DCP1 and DCP2 proteins, the plasmids expressing DCP1 and DCP2 were co-transfected into insect cells, followed by Co-IP using DCP2-specific antibody. Western blots revealed that the DCP1 protein was directly interacted with the DCP2 protein (Fig 2B).

**Fig 2.**
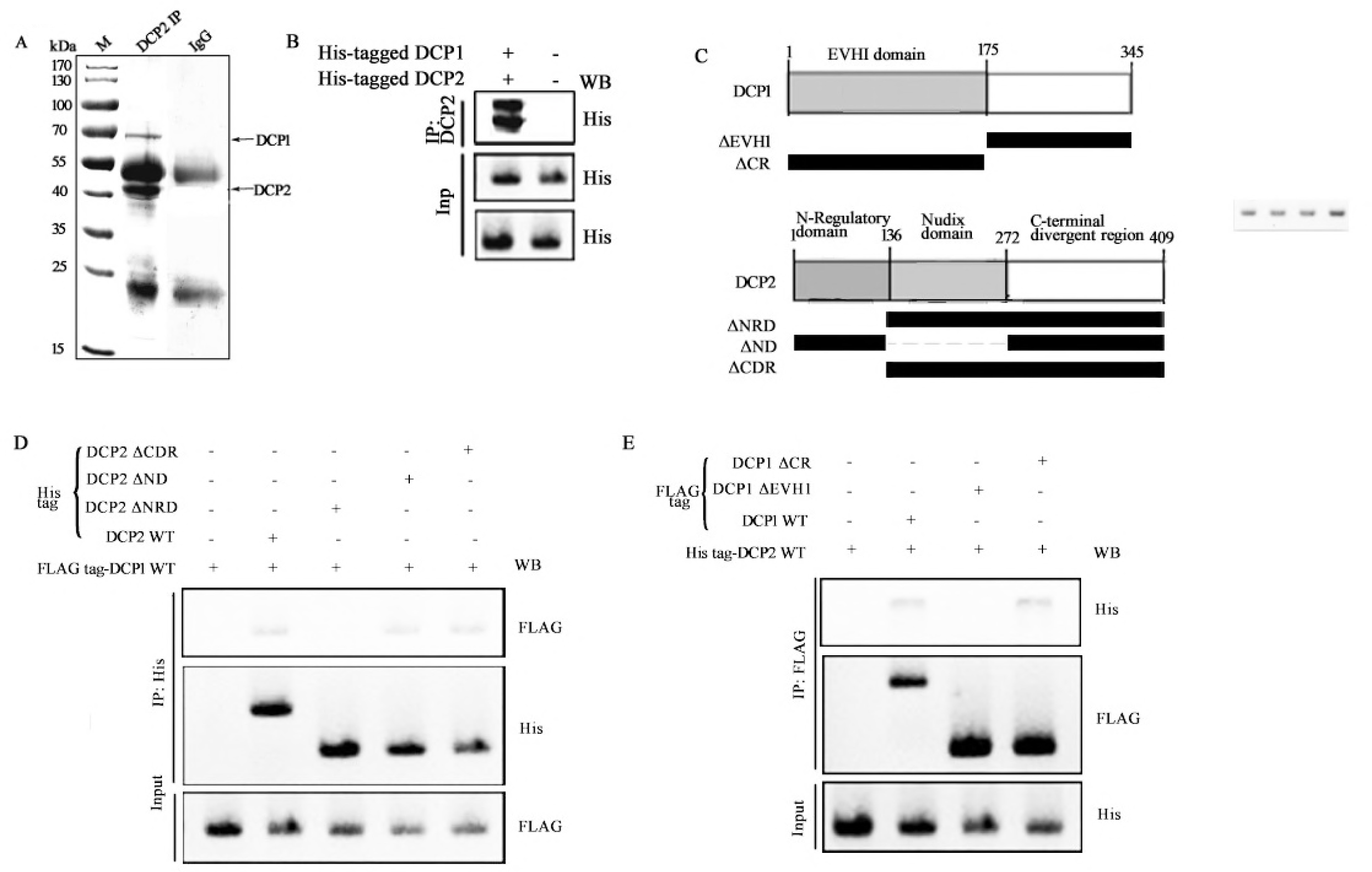
Proteins interacted with DCP2. (A) The proteins bound to DCP2. Co-IP using the DCP2-specific antibody was conducted. The eluted proteins were subjected to SDS-PAGE, followed by protein identification using mass spectrometry. (B) The interaction between DCP1 and DCP2 proteins. The His-tagged DCP1 and DCP2 were co-transfected into insect cells. At 48 h after co-transfection, Co-IP was conducted using DCP2-specific antibody, followed by Western blot analysis with anti-His IgG. (C) The constructs of DCP1 and DCP2 domain deletion mutants. (D) and (E) The interactions between DCP1 and DCP2 domains. The full-length and/or deletion mutants of DCP1 and DCP2 were co-transfected into insect cells. At 48 h after transfection, the target proteins were immunoprecipitated with anti-His (D) or anti-FLAG IgG (E), followed by Western blot analysis.

To identify which domains of DCP1 and DCP2 were interacted, the deletion mutants of DCP1 EVH1 domain (ΔEVH1, FLAG-tagged), DCP1 C-terminal region (ΔCR, FLAG-tagged), DCP2 N-terminal regulatory domain (ΔNRD, His-tagged), DCP2 Nudix domain (ΔND, His-tagged) and DCP2 C-terminal divergent region (ΔCDR, His-tagged) were constructed, respectively (Fig 2C). The deletion constructs and the full-length DCP1 (FLAG-tagged) or DCP2 (His-tagged) were co-transfected into insect cells. The results showed that when insect cells were co-transfected with DCP2 ΔNRD and full-length DCP1, the DCP1 protein was not detected in the immunoprecipitated product using His antibody (Fig 2D), showing that DCP1 was interacted with DCP2 N-terminal regulatory domain. When DCP2 and DCP1 ΔEVH1 were co-transfected into cells, the DCP2 protein did not exist in the immunoprecipitated complex (Fig 2E), indicating that DCP2 was interacted with DCP1 by binding to its EVH1 domain.

The above findings indicated that the EVH1 domain of DCP1 was interacted with the N-terminal regulatory domain of DCP2.

### Role of shrimp DCP1 in virus infection

To explore the influence of DCP1 on virus infection of shrimp, the expression profile of DCP1 was examined in hemocytes of WSSV-infected shrimp. The data of Northern blots and Western blots indicated that the DCP1 expression was significantly upregulated in shrimp in response to WSSV infection, suggesting the involvement of DCP1 in virus infection (Fig 3A).

**Fig 3.**
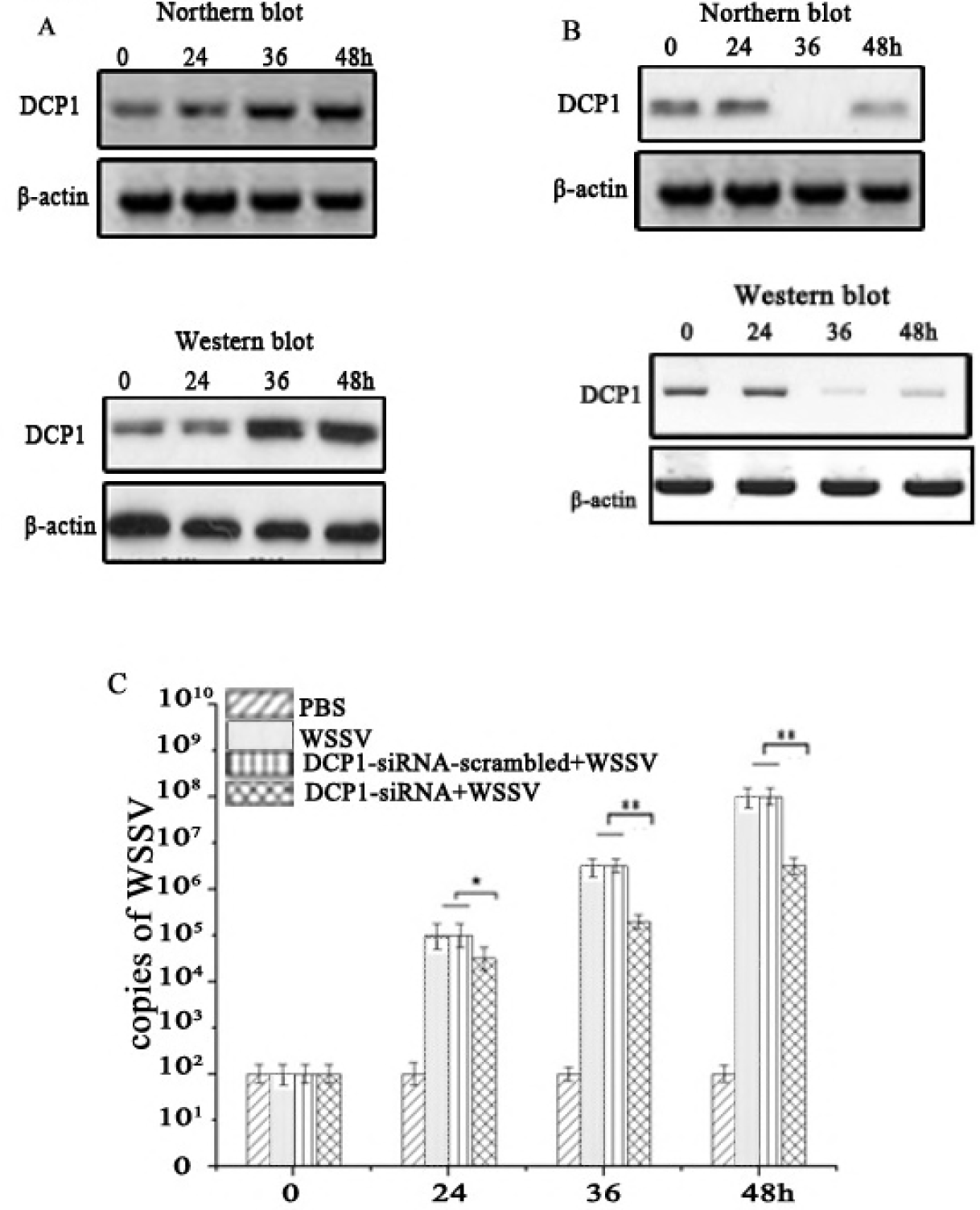
Role of shrimp DCP1 in virus infection. (A) DCP1 expression profile in shrimp in response to virus infection. Shrimp were challenged with WSSV. At different times post-infection, the expression of DCP1 was examined in shrimp hemocytes by Northern blotting or Western blotting. Shrimp β-actin was used as a control. The numbers indicated the time post-infection. Probes or antibodies used were shown on the left. (B) Silencing of DCP1 in shrimp. Shrimp were injected with DCP1-siRNA, followed by the detection of DCP1 with Northern blot or Western blot. The probes or antibodies were indicated on the left. (C) Influence of DCP1 silencing on virus infection. WSSV and DCP1-siRNA or DCP1-siRNA-scrambled were co-injected into shrimp. WSSV alone and PBS were used as controls. At different time after injection, the WSSV copies in shrimp were examined with quantitative real-time PCR (*, *p*<0.05; **, *p*<0.01).

In an attempt to assess the role of DCP1 in virus infection, the DCP1 expression was knocked down by sequence-specific siRNA in WSSV-infected shrimp, followed by evaluation of virus infection. The results revealed that the expression of DCP1 was silenced by DCP1-siRNA (Fig 3B). The DCP1 silencing led to significant decreases of WSSV copies compared with the controls (WSSV and WSSV+DCP1-siRNA-scrambled) (Fig 3C). These findings indicated that DCP1 played a positive role in virus infection.

### Effects of the interaction between shrimp miR-87 and *DCP2* on virus infection

To reveal the miRNAs targeting shrimp *DCP2* gene, the miRNAs targeting *DCP2* were predicted. The prediction data showed that the shrimp *DCP2* gene was a potential target of miR-87 (Fig 4A). To evaluate the interaction between miR-87 and *DCP2* gene, the plasmid pIZ/EGFP-DCP2-3’UTR containing the EGFP and the DCP2 3’UTR was co-transfected with miR-87 into the insect cells. The results indicated that the fluorescence intensity of the cells co-transfected with miR-87 and pIZ/EGFP-DCP2-3’UTR was significantly decreased compared with the controls (Fig 4B). However, the fluorescence intensity of the cells co-transfected with miR-87 and EGFP-ΔDCP2-3’UTR was similar to those of the controls (Fig 4B). These findings revealed that miR-87 was directly interacted with *DCP2* gene. In order to examine the interaction between miR-87 and *DCP2* gene *in vivo*, miR-87 was overexpressed in shrimp, followed by the analysis of *DCP2* gene expression. It was revealed that the miR-87 overexpression led to a significant decrease of *DCP2* expression at transcript and protein levels compared with the controls (Fig 4C), indicating that miR-87 was interacted with *DCP2* gene *in vivo*.

**Fig 4.**
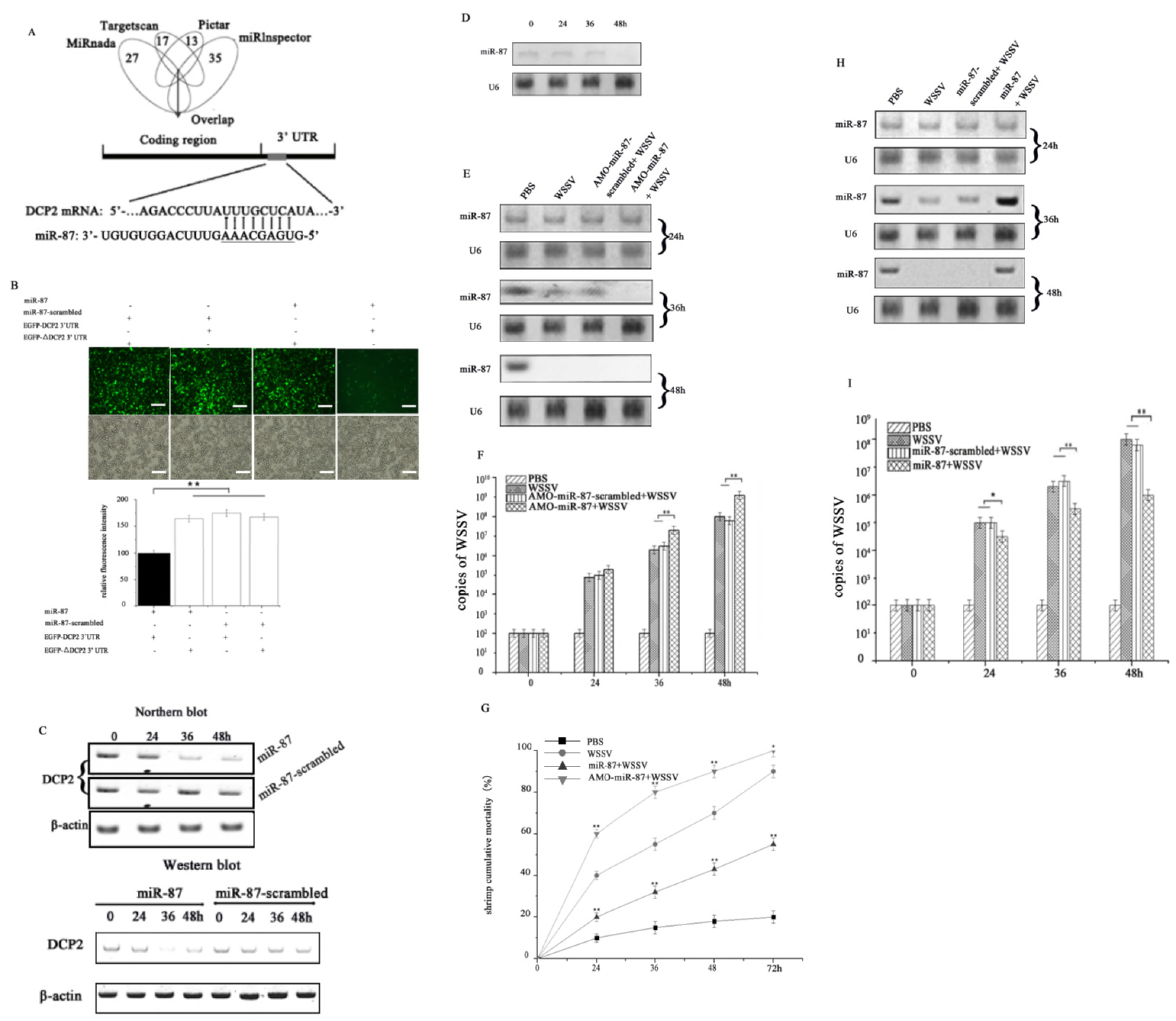
Effects of the interaction between shrimp miR-87 and *DCP2* on virus infection. (A) Prediction of miRNAs targeting *DCP2*. According to the prediction, the 3’ UTR of *DCP2* gene could be targeted by miR-87. The seed sequence of miR-87 was underlined. (B) Direct interaction between miR-87 and DCP2 3’ UTR. The insect High Five cells were co-transfected with miR-87 and the plasmid EGFP-DCP2-3’UTR or EGFP-ΔDCP2-3’UTR. At 36 h after co-transfection, the fluorescence intensity of insect cells was evaluated. Scale bar, 50 μm. (C) Interaction between miR-87 and DCP2 *in vivo*. MiR-87 was overexpressed in shrimp. At different time after miR-87 overexpression, the *DCP2* mRNA and protein levels were examined by Northern blot and Western blot, respectively. As a control, miR-87-scrambled was included in the assays. Data were representatives of three independent experiments. The probes or antibodies were indicated on the left. (D) Expression level of miR-87 in virus-infected shrimp. Shrimp were challenged with WSSV. At different time post-infection, miR-87 was detected in hemocytes of virus-infected shrimp by Northern blotting. U6 was used as a control. Probes were indicated on the left. (E) Silencing of miR-87 expression in shrimp. Shrimp were co-injected with AMO-miR-87 and WSSV. As a control, AMO-miR-87-scrambled was included in the injection. At different time post-infection, miR-87 was detected by Northern blot. The probes used were indicated on the left. The numbers showed the time points post-infection. U6 was used as a control. (F) Influence of miR-87 silencing on virus copies. WSSV and AMO-miR-87 or AMO-miR-87-scrambled were co-injected into shrimp. WSSV and PBS were used as controls. At different time after injection, the WSSV copies in shrimp were examined with quantitative real-time PCR. (G) Effects of miR-87 silencing or overexpression on WSSV-infected shrimp mortality. (H) Overexpression of miR-87 in shrimp. Shrimp were co-injected with miR-87 or miR-87-scrambled and WSSV. At different time after injection, the shrimp were subjected to Northern blot with probes indicated on the left. PBS and WSSV were used as controls. (I) Impact of miR-87 overexpression on WSSV copies. Shrimp were simultaneously injected with miR-87 and WSSV. As a control, miR-87-scrambled was included in the injection. At different time post-infection, the virus copies were examined with quantitative real-time PCR. In all panels, the significant differences between treatments were indicated (*, *p*< 0.05; **, *p*< 0.01).

To explore the role of shrimp miR-87 in virus infection of shrimp, the expression level of miR-87 was examined in hemocytes of WSSV-infected shrimp. Northern blots indicated that the host miR-87 expression was significantly downregulated in shrimp in response to WSSV infection, suggesting that miR-87 played an important role in the shrimp antiviral immunity (Fig 4D).

In order to assess the influence of miR-87 on virus infection, the miR-87 expression was silenced or overexpressed in the WSSV-infected shrimp, followed by the evaluation of virus infection. The results showed that the expression of miR-87 was knocked down by AMO-miR-87 compared with the controls (Fig 4E). The miR-87 silencing led to significant increases of WSSV copies and the virus-infected shrimp mortality compared with the controls (Fig 4F and 4G). On the other hand, when miR-87 was overexpressed (Fig4H), the WSSV copies and the virus-infected shrimp mortality were significantly decreased compared with the controls (Fig 4G and 4I).

Taken the above data together, these findings presented that miR-87 could inhibit virus infection in shrimp by targeting shrimp *DCP2* gene.

### Influence of viral WSSV-miR-N46 targeting *DCP1* on virus infection

To characterize the miRNAs targeting *DCP1*, the viral miRNAs targeting *DCP1* gene were predicted. The miRNA target prediction showed that the *DCP1* gene might be the target of WSSV-miR-N46, a viral miRNA encoded by WSSV (Fig 5A). To validate the target prediction, the synthesized viral miRNA and the plasmid EGFP-DCP1-3’ UTR were co-transfected into insect cells. The results indicated that the fluorescence intensity of the cells co-transfected with WSSV-miR-N46 and EGFP-DCP1-3’ UTR was significantly decreased compared with that in the controls (Fig 5B), showing that WSSV-miR-N46 was directly interacted with *DCP1* gene.

**Fig 5.**
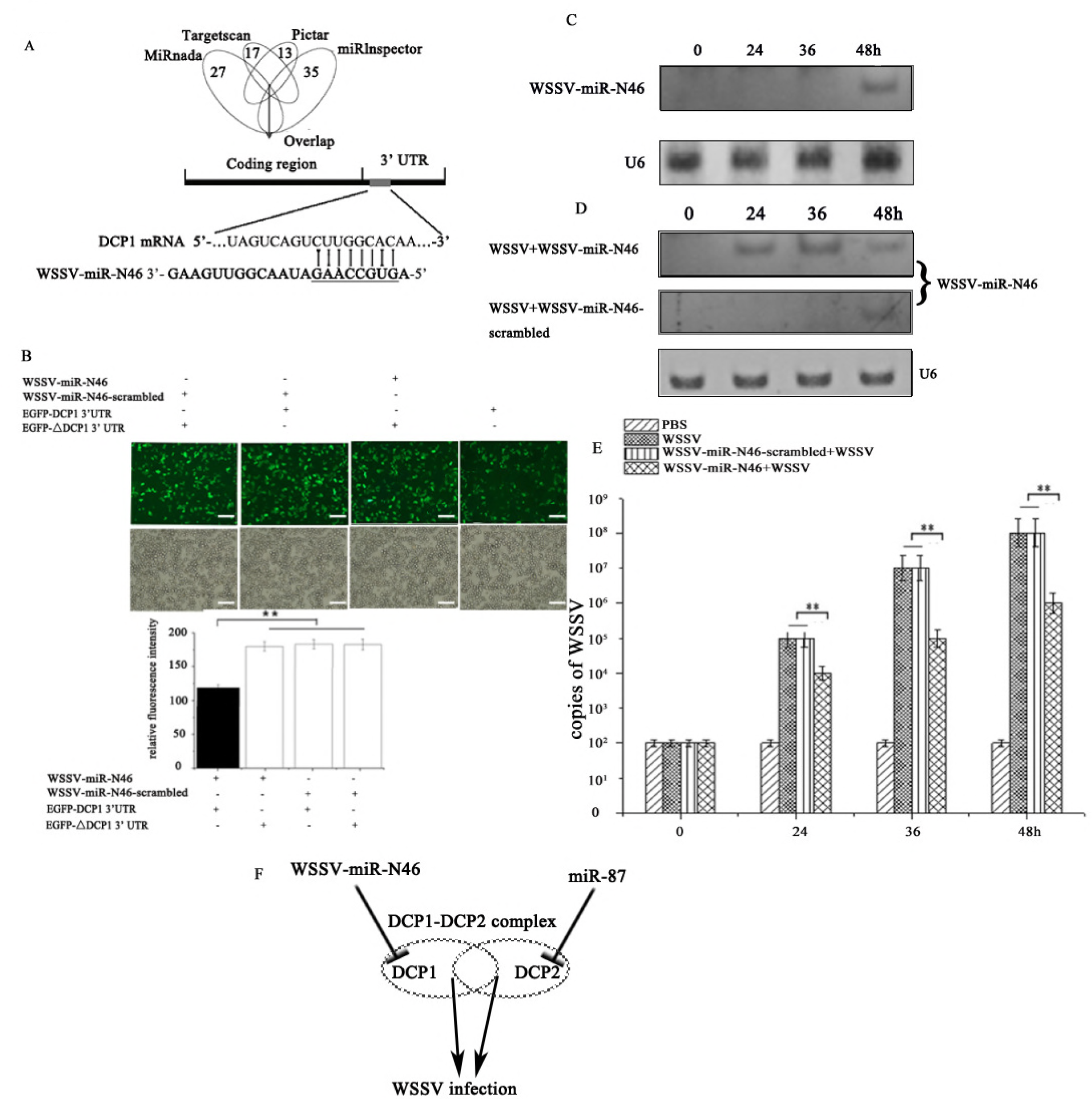
Influence of viral WSSV-miR-N46 targeting *DCP1* on virus infection. (A) The prediction of viral miRNA targeting *DCP1*. As predicted, the 3’ UTR of *DCP1* was targeted by WSSV-miR-N46, a WSSV-encoded viral miRNA. The seed sequence was underlined. (B) The direct interaction between WSSV-miR-N46 and *DCP1* gene in insect cells. Insect High Five cells were co-transfected with WSSV-miR-N46 or WSSV-miR-N46-scrambled and EGFP, EGFP-DCP1 3’ UTR or EGFP-ΔDCP1 3’ UTR. At 48 h after co-transfection, the fluorescence of cells was examined (**, *p*<0.01). Scale bar, 50 μm. (C) The expression pattern of WSSV-miR-N46 in shrimp in response to virus infection. Shrimp were challenged with WSSV. At different time post-infection, WSSV-miR-N46 was detected by Northern blotting. U6 was used as a control. The number indicated the time points post-infection. Probes were indicated on the left. (D) The overexpression of WSSV-miR-N46 in shrimp. Shrimp were simultaneously injected with WSSV and WSSV-miR-N46. As a control, WSSV-miR-N46-scrambled was included in the injection. At different time post-infection, shrimp hemolymph was subjected to Northern blotting. U6 was used as a control. The probes were shown on the right. (E) The influence of WSSV-miR-N46 overexpression on WSSV infection. Shrimp were simultaneously injected with WSSV-miR-N46 and WSSV. As a control, WSSV-miR-N46-scrambled was included in the injection. At different time post-infection, the WSSV copies were examined with quantitative real-time PCR (**, *p*<0.01). (F) Mode for the miRNA-mediated signaling pathway in virus infection.

In an attempt to reveal the role of WSSV-miR-N46 in virus infection, the expression of WSSV-miR-N46 in WSSV-challenged shrimp was examined. Northern blotting results indicated that WSSV-miR-N46 was detected at 48 h after virus infection in shrimp (Fig 5C). Therefore, WSSV-miR-N46 was overexpressed in shrimp (Fig 5D), followed by evaluation of virus copy. The results revealed that the WSSV-miR-N46 overexpression significantly decreased the number of WSSV copies in shrimp (Fig 5E), indicating that WSSV-miR-N46 played a negative role in WSSV replication.

Taken together, the findings revealed that the viral miRNA (WSSV-miR-N46) and host miRNA (miR-87) suppressed virus infection by targeting the DCP1-DCP2 complex (Fig 5F).

## Discussion

As reported, the DCP1-DCP2 complex, localized in processing bodies (P bodies), can regulate the animal antiviral immunity by two strategies, that is the decapping of retrovirus RNAs and the suppression of RNAi pathway (32–36). During the process of retrovirus infection, the canonical mRNA decapping enzyme DCP2, along with its activator DCP1, could restrict the infection of retrovirus at the level of mRNA transcription (34, 35). The host DCP1-DCP2 complex directly decapps retrovirus mRNAs or cellular mRNAs targeted by bunyaviruses for cap-snatching, thus creating a bottleneck for retrovirus replication (33, 35). During the infection of Sindbis virus or Venezuelan equine encephalitis virus, the host can inhibit the infection of retrovirus through the DCP1-DCP2-mediated 5’-3’ decay pathway. During the bunyaviruses infection in the insects and mammals, the bunyaviruses cap their mRNAs at the 5’ ends by the “cap-snatching” machinery in the P bodies (35). The virally encoded nucleocapsid N protein recognize 5’ caps and 10-18 nucleotides (nt) downstream 5’ caps of cellular mRNAs and the viral RNA-dependent RNA polymerase cleaves the mRNA at the same position. Subsequently the cleaved 5’ caps are used for viral mRNA synthesis (35). Regarding the role of the DCP2-DCP2 complex in RNAi pathway, it is found that the silencing of DCP2 and/or DCP1 promotes RNAi, showing that the DCP2-DCP1 complex takes a negative effect on the RNAi pathway (35, 36). RNAi, an important component of innate immune responses, mediated by siRNAs or miRNAs, plays crucial roles against virus infection in invertebrates and plants that rely solely on innate mechanisms to combat viral infection (30, 34, 37). Up to date, however, little is known about the role of the DCP1-DCP2 complex in DNA virus infection. In the present study, the findings indicated that the silencing of the DCP1-DCP2 complex inhibited the infection of WSSV, a DNA virus of shrimp, suggesting that the DCP1-DCP2 complex facilitated DNA virus infection. Due to the suppressive role of the DCP1-DCP2 complex in RNAi pathway against virus infection (35, 36), the DCP1-DCP2 complex could promote WSSV infection in shrimp. In this context, our study contributed a novel aspect of the DCP1-DCP2 complex in virus-host interactions.

In the present investigation, the results showed that the host and viral miRNAs could inhibit the expressions of DCP1 and DCP2 during DNA virus infection. MiRNAs, a large class of small noncoding RNAs in diverse eukaryotic organisms, are sequentially processed by two RNase III proteins, Drosha and Dicer from the stem regions of long hairpin transcripts (28, 37). The mature miRNA strand is liberated from the miRNA:miRNA* duplex and integrated into the RNA induced silencing complex (RISC), and inhibits the expression of cognate mRNA through degradation or translation repression in the RISC (18). During virus infection the host miRNAs or/and viral miRNAs can regulate virus infection by targeting viral or/and host genes (2, 17, 23, 24, 27–29, 31, 32, 38). As well reported, the virus-encoded miRNAs (viral mRNAs) can target virus and/or host genes, leading to virus infection or virus latency (29, 32, 38, 39). In shrimp, a viral miRNA WSSV-miR-N12 targets the virus wsv399 gene, resulting in virus latency (32). The viral miRNA-mediated regulation of virus infection or virus latency is an efficient strategy for virus to escape its host immune responses. However, the involvement of miRNA in the degradation of cellular mRNAs mediated by DCP1-DCP2 complex has not been explored. Our study revealed that the host and viral miRNAs could regulate the DCP1-DCP2 complex to affect virus infection. Therefore, our study provided novel insights into the regulatory mechanism of DCP1-DCP2 complex in virus-host interactions and that the miRNA-mediated regulation of DCP1-DCP2 complex took great effects on RNAi immunity of invertebrates against virus infection.

## Acknowledgements

This work was supported by National Natural Science Foundation of China (31430089) and National Program on the Key Basic Research Project (2015CB755903).

